# Changing variability is an overlooked aspect of protected area planning

**DOI:** 10.1101/2023.10.26.564062

**Authors:** Rekha Marcus, Michael J. Noonan

**Affiliations:** Department of Biology, The University of British Columbia Okanagan, Kelowna, British Columbia, Canada; Okanagan Institute for Biodiversity, Resilience, and Ecosystem Services, The University of British Columbia Okanagan, Kelowna, British Columbia, Canada; Department of Computer Science, Math, Physics, and Statistics, The University of British Columbia Okanagan, Kelowna, British Columbia, Canada

**Keywords:** climate change, conservation planning, priority conservation areas, reserve design, stochasticity, variance

## Abstract

Protected areas are widely used management tools designed to support the long-term conservation of biodiversity. The effectiveness of protected areas is being challenged by human-induced climate change, however, which is causing three broad shifts away from the current distribution of climate trends: a change in mean conditions, a change in the variance around the mean, and/or a change in symmetry. Though changes in average conditions are certainly important, the second behaviour, a change in variance, brings a unique set of challenges that species must respond to. As conditions become more variable, phenological events become less predictable, extreme weather events become more frequent, food security and ecosystem stability are compromised, and extinction risk increases. It therefore stands to reason that changes in the variance of local conditions should be a core consideration when designing protected areas.

Here, we reviewed the literature to determine the extent to which changes in variance are being incorporated into protected area planning. Worryingly, we found that fewer than a quarter of the 100 studies we surveyed formally considered how climate change might change mean conditions, and only four considered climate change-induced changes in the variance around the mean. Our evaluation reveals an alarming gap in protected area research. The majority of researchers continue to make recommendations for protected areas without acknowledging that the area(s) they are recommending for protection may have markedly different conditions in the future. Whether variability is considered or not, stochastic events represent a serious threat to the persistence of species and complex ecosystems. Effective conservation requires actively considering how the stability of conditions within protected areas will be impacted by future climate change. As global climate patterns tend towards increasing unpredictability, protecting less variable habitat should be a priority to ensure local populations are not exposed to elevated extinction risk.

## Introduction

Protected areas (PAs) are ecosystem-scale management tools aimed at mitigating the impacts of anthropogenic disturbance on the natural world and preserving culturally and ecologically important landscapes (Le Saout et al., 2013; Naughton-Treves et al., 2005). Though aspects of PA creation and management are debated (e.g., Brockington, 2004; Oldekop et al., 2016), their conservation value is well recognized (Chen et al., 2022; Nagendra, 2008; Rodrigues et al., 2004) and they remain widely used management tools. Yet, for PAs to effectively achieve their ultimate goal of protecting biodiversity and ecosystem function, it is essential that they be placed in areas where they will have maximal benefit (Le Saout et al., 2013). To this end, the International Union for Conservation of Nature (IUCN) recommends that PAs should be created “*to achieve the long term conservation of nature with associated ecosystem services and cultural values*” (IUCN 2008). While ‘long term conservation’ is at the heart of the PA concept, this goal is being challenged by human-induced climate change (Field et al., 2012), with some estimates suggesting that as few as 8% of protected areas will remain effective by the end of the century (Loarie et al., 2009). It therefore stands to reason that the impacts of climate change on natural ecosystems should be core considerations for the creation of PAs that will safeguard biodiversity for generations to come (Gillingham et al., 2015; Hannah et al., 2007).

From the perspective of PA planning, climate change is expected to result in three broad shifts away from the current distribution of climate trends: a change in the mean, a change in the variance, and/or a change in symmetry (formally the first, second, and third statistical moments; IPCC, 2021; Field et al., 2012; Parey et al., 2010). Though changes in average conditions can certainly impact the effectiveness of PAs (Loarie et al., 2009), the second behaviour, a change in variance, brings a unique set of challenges that species must respond to. As local conditions become more variable, phenological events become less predictable (Menzel et al., 2006), the frequency of extreme weather events increases (Stott, 2016; Trenberth et al., 2015), food security and ecosystem stability become compromised (Thornton et al., 2014; White et al., 2021), and extinction risk increases (Easterling et al., 2000; Vincenzi, 2014). As negative ecological responses to extreme weather events have been recorded in 57% of all species (Maxwell et al., 2019), habitats that are expected to remain stable under future climate change should be prioritized as PAs. Yet, while mentions of less predictable weather and extreme events are becoming more frequent in public discourse and conservation research alike (Maxwell et al., 2019; Silver & Andrey, 2019), the extent to which variance in local conditions is actively incorporated into PA research remains limited. Here, we review the peer-reviewed literature to explore the extent to which researchers are considering changing variability in conditions when attempting to identify priority areas. Our evaluation reveals a worrying gap in PA research, and provides essential context for future research aimed at improving PA creation and management in the face of human-induced climate change.

## Methods

Our aim was to determine trends in the information being used to identify priority conservation areas, whether by statistical analysis, a literature review, or employing a theoretical framework. Using both Web of Science and Google Scholar, studies were identified via the search terms “conservation areas”, “reserve design”, “conservation planning”, “site selection”, and “priority conservation areas”. The Web of Science search was last accessed on May 31, 2023, and the Google Scholar search was last accessed September 18, 2023. No additional filters were added to the search, and results were sorted by most relevant in both search engines. From these lists, the first 1000 studies generated from both websites were subset and the first 100 most relevant studies were selected. The most relevant papers were ones that specifically addressed the creation of protected areas. Any studies focused on anthropogenic and/or socio-economic responses to PAs or PA management were excluded. Each paper was first classified according to how it considered climate change, namely: i) made no mention of change, ii) mentioned a change in mean conditions but did not include this in the analyses, iii) incorporated a change in mean conditions in their analysis, iv) mentioned a change in variance but did not include this in the analyses, and v) incorporated a change in variance in their analysis. In addition, we recorded the year of publication, journal impact factor, geographic extent, the number and types of measures used for the analysis, the study methods, whether the study concluded in a specific recommendation for a PA to be created, as well as data availability. Because authors made use of many different variables in their analyses, we classified these as being either static (i.e., variables that do not change over short time periods, such as elevation or species endemism indices), or dynamic (i.e., variables that have the capacity to change, such as temperature or environmental productivity) and recorded the use of these two variable classes. As a note, the majority of biodiversity measures (e.g., species distributions) were considered to be static; although these do have the ability to change over short periods of time, the majority of the biodiversity measures in the studies were taken over a short time span and thus could not be used to determine how they might change over time. Variables that were unclear as to whether they were static or dynamic were considered to be static unless the subsequent analysis made explicit mention of changes in conditions, at which point these variables were then considered dynamic. Variables were then further categorized as being measures of i) biodiversity, ii) geophysical landscape, iii) human use, iv) meteorology, and v) productivity. Some studies did not make use of any variables or mathematical analysis, in which case these categories of the data were simply recorded as N/A.

The full dataset and paper lists from the search results are provided in the Supplementary Material (S1, S2, and S3, respectively). Though our literature search was not exhaustive, our compiled dataset is likely to be representative of broad trends in the field. Figures and descriptive statistics were generated in R (Ver. 4.3.1, R core team), and all of the code and data are openly available at https://github.com/QuantitativeEcologyLab/variance-parks.

## Results

The 100 studies we surveyed were published between the years 1998 and 2023 (Fig. 1A), and were relatively evenly distributed across the globe, though Asia was the most heavily represented (Fig. 1B). We noticed a wide range methodological approaches, with 72% of studies making use of statistical or mathematical modelling in order to determine priority conservation areas, and the remainder presenting theoretical approaches to PA planning (25%) or literature reviews of specific approaches (3%).

**Figure 1:**
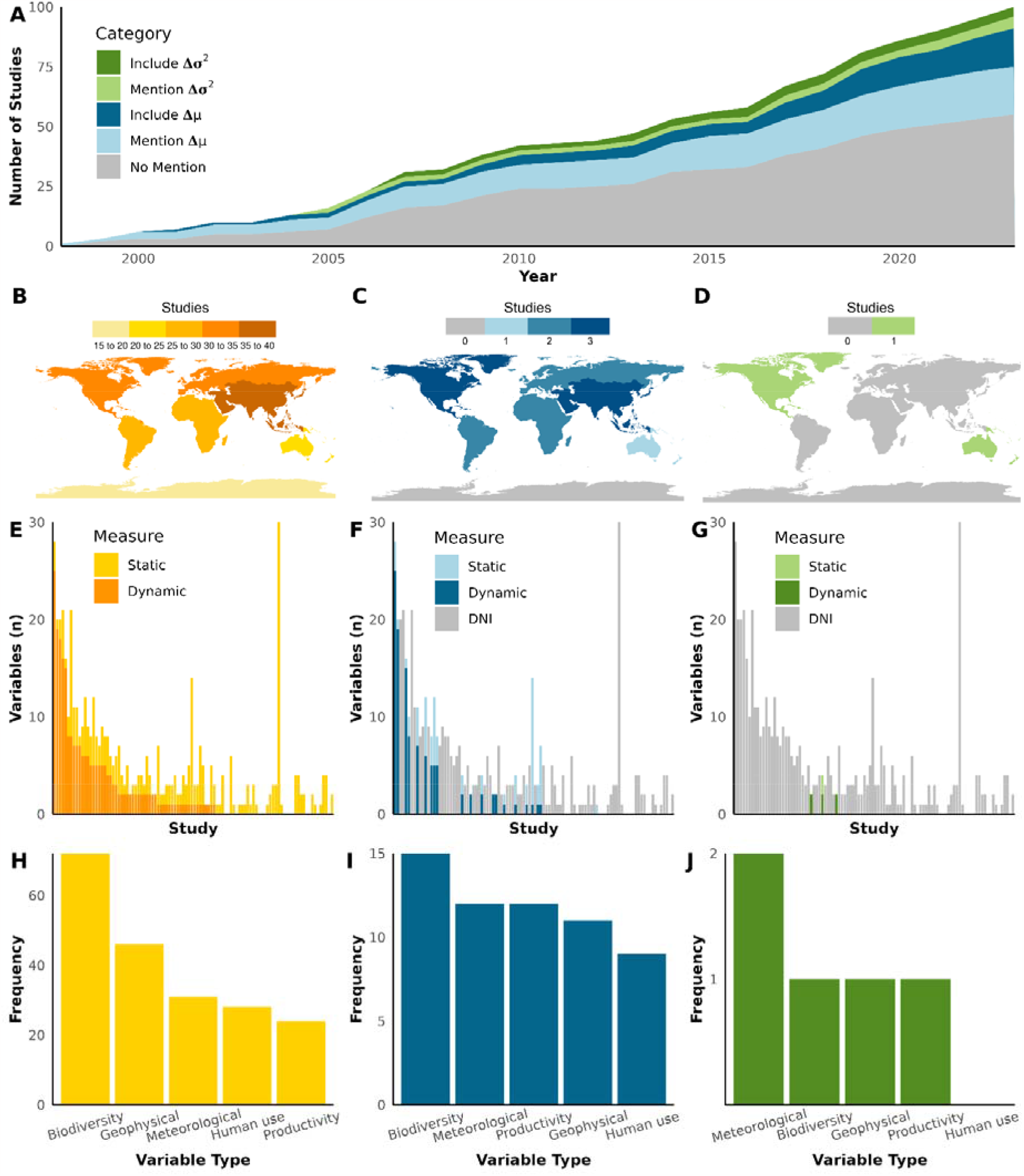
Descriptive statistics of studies on protected area planning, subset by all studies (yellow, n = 100), studies that incorporated changes in mean (blue, n = 24), and studies that incorporated changes in variance (green, n = 4). Panel (A) shows the cumulative number of studies that either i) made no mention of change (no mention), ii) mentioned a change in mean conditions but did not include this in the analyses (mention Δμ), iii) incorporated a change in mean conditions in their analysis (include Δμ), iv) mentioned a change in variance but did not include this in the analyses (mention Δσ^2^), or v) incorporated a change in variance in their analysis (include Δσ^2^). The second row depicts the geographic distribution of (B) all studies, (C) studies that included changes in mean, and (D) studies that included changes in variance, with darker shading indicating a higher number of studies. Map lines delineate study areas and do not necessarily depict accepted national boundaries. Note: studies that did not focus on specific geographic areas are not shown. Panels (E), (F), and (G) show the number of static versus dynamic variables use in each study, ordered by their use of dynamic variables. Panels (H), (I), and (J) depict the distribution of variables used in each category to quantify habitat quality.

The use of static versus dynamic variables was mixed, though static variables were the most commonly used, with studies using an average of 1.5 static variables for every dynamic variable. This ranged substantially though, with 37 studies relying solely on static variables with no capacity to model change, and 5 studies relying entirely on dynamic variables (Fig. 1E). The most commonly used variables for identifying priority areas were measures of biodiversity, typically species distributions (Fig. 1H). Nonetheless, measures describing the geophysical landscape, the amount of human use, meteorology, and productivity were also regularly employed (Fig. 1H).

### Few studies accounted for changing mean conditions

Surprisingly, 55% of studies made no mention of climate change, nor how environmental conditions might change in the future. Further, of the 45 studies that did mention changes in mean conditions, many failed to actually account for this in their analyses. As such, only 24% of studies formally considered potential changes in mean conditions over time in the regions they assessed. The geographic extent of studies that incorporated changing mean conditions largely mirrored the overall distribution of priority area research (Fig. 1C). Studies in this category typically projected future conditions based on past data and various climate change-based prediction models, primarily via measures of biodiversity (n = 15), meteorology (n = 12) and productivity (n = 12; Fig 1I). Notably, these studies all made use of dynamic measures in order to understand how these locations would change over time (Figure 1F).

### Fewer studies accounted for changing variance

Only four of the 100 studies we evaluated incorporated changing variance into their analyses. These were limited to North America and Oceania (n = 1 for each continent; Fig. 1D), as well as two theoretical studies that did not focus on any specific area. These four studies relied primarily on dynamic variables, modelling patterns in mean and variance over time and recommended priority conservation areas based on the threat that changing variances and extreme events represented. These were most commonly meteorological measures such as annual temperature and precipitation trends, but also included elevation and species-specific habitat use, such as nesting areas (Fig. 1G). Interestingly, studies in this category did not include variables related to human use, such as the human footprint index or road densities.

## Discussion

Researchers regularly make recommendations on the creation of protected areas. The analyses supporting these recommendations typically involve surveying (an) aspect(s) of the land and the biodiversity within it, and then leveraging this information to make policy recommendations. Worryingly, however, fewer than a quarter of the studies we surveyed formally considered how climate change might change mean conditions, and fewer still considered climate change-induced changes in variance. Whether variability is considered or not, stochastic events represent a serious threat to the persistence of species and complex ecosystems (Easterling et al., 2000; Pressey et al., 2007; Vincenzi, 2014). The studies we reviewed were published between 1998 and 2023, a timeframe during which climate change and extreme weather events would reasonably be expected to be a consideration in priority area research. Yet, we found that the majority of researchers (72% of the studied literature) continue to make recommendations for protected areas without acknowledging that the area(s) they are recommending for protection may have markedly different conditions in the future.

Effective conservation requires actively considering how the stability of protected areas will be impacted by future climate change (Gillingham et al., 2015; Hannah et al., 2007). As global climate patterns tend towards increasing unpredictability, protecting less variable habitat should be a priority in order to ensure local populations are not exposed to elevated extinction risk (Easterling et al., 2000; Vincenzi, 2014). Despite the widespread attention that climate change has gained in recent years (Silver & Andrey, 2019), however, our work highlights how the consideration of changing variability in local conditions still represents a major gap in protected area planning. Though it was surprising to see that so few studies accounted for changing variability in their analyses, this finding mirrors a review of climate change studies by Thompson et al. (2013), who found only 6.1% of terrestrial studies considered the extreme events caused by climate change – the majority of the studies only included mean temperature increases, and did not consider variability or extreme events at all. While it is the norm, continuing to ignoring variance is a dangerous omission, as some significant weather shifts can occur that generate only a small change in mean conditions, but substantially increase the variance (Mearns et al., 1984). Thus, albeit some scientists having identified the need to include measures of change in variance in conservation planning (Lewis & King, 2017; Mearns et al., 1984; Pressey et al., 2007; Stouffer & Wetherald, 2007; Thompson et al., 2013), our review highlights that this has yet to be broadly reflected in the literature.

Though the lack of research on changing variance was worrying, a number of authors did actively consider changing stochasticity when attempting to identify priority conservation areas. In this regard, the four studies we reviewed that did account for a change in variance incorporated dynamic variables in their analyses. For example, Butt et al. (2016) presented recommendations for priority conservation areas for sea turtle nesting sites based on an analysis that explored how sea level rise would increase the probability of flooding events. Similarly, Leroux et al. (2007)proposed the concept of minimum dynamic reserves, which take into account the rate of natural disturbances and the effects these have on ecological communities. The work of these authors highlights the importance of leveraging the information contained in dynamic variables when attempting to identify priority conservation areas. With this in mind, we did find that the majority of studies included at least one dynamic variable in their analyses, which is a prerequisite for being able to consider changing variance. However, more than 1/3 of studies did rely entirely on the use of static variables, which, while not without value, provide no capacity to model how conditions will change over time. An exception of note to the use of species distributions as a static measure was a study conducted by Fogarty et al. (2020), which studied changing forest sizes, grassland cover, and woody plant encroachment. In order to consider the effects of climate change in PA planning, we recommend that researchers make greater use of dynamic variables, taking into account the timescales over which these can change, when identifying priority conservation areas. This is especially important when considering species distributions and other biodiversity measures, as climate change can cause species distributions can change substantially over time (Loarie et al., 2009), especially in response to evolving threats and extreme events (Stanton et al., 2012).While modelling variances represents a technical challenge, it is essential for effective conservation (Pressey et al., 2007). Though we found that many studies relied on relatively inflexible conservation planning software such as MARXAN or MAXENT in their analyses, the increasing availability of open-source data and more flexible analytical tools should continue to make such analyses more accessible.

While not a primary focus of our review, we also noted many shortcomings in terms of accessibility and reproducibility. In particular, only 14% of the studies reviewed included specific data on the areas they were recommending be protected, usually in the form of spatial coordinates or a GIS shapefile (Belote & Wilson, 2020; Sutton et al., 2023). Notably, none of the 4 studies that accounted for changes in variance had data from their recommendations readily available. The overwhelming majority of papers thus provided only the content of their papers as material for policymakers to work with. In the era of Open Science, where the field should be trending towards increased transparency and availability of data, this was troubling to observe. Beyond this lack of transparency, many of the studies used complex language that could be challenging to understand and interpret. Thus, despite the fact that studies may have made valuable contributions to PA research, due to a tendency for poor data management and accessibility practices, many PAs risk not yielding concrete conservation action. We therefore recommend that conservation scientist make improving access to their underlying data, models, and results a priority going forward (see also Roche et al., 2022).

## Conclusions

Protected areas continue to be one of the primary tools for safeguarding biodiversity. Indeed, this was highlighted at the recent COP15 meeting, at which many states of the world agreed on a historic package to protect 30% of the world’s land mass by 2030 (i.e., the 30 by 30 initiative; Kunming-Montreal Global Biodiversity Framework, 2022). With only ∼16% of the world’s land area currently lying within PAs, achieving this goal will require protecting an additional ∼2.8 million ha of habitat within the next ca. 6 years (Zeng et al., 2022). At the cusp of this global effort towards the expansion of protected areas, the results of our study highlight a dangerous gap in protected area research the risks challenging their effectiveness. The lack of studies modelling environmental variability and accounting for how this may change suggests that many protected areas might be vulnerable to climate change-induced increases in stochastic weather events (Loarie et al., 2009). To ensure the long-term protection of ecologically and culturally valuable biodiversity, current approaches to protected area research must change. By filling in the gaps of PA networks with robust regions that can provide stable conditions and resources, we can take a valuable step towards safeguarding the planet’s biodiversity.

## Acknowledgments and data

We thank the members of the Quantitative Ecology Lab for their feedback on early drafts of the manuscript. This work was supported by the NSERC Discovery Grant RGPIN-2021-02758, the Canadian Foundation for Innovation, as well as the University of British Columbia, Okanagan. The compiled dataset can be found in the supplementary information. All data and code used to generate the figures can be found in the GitHub repository at https://github.com/QuantitativeEcologyLab/variance-parks.

## Supplementary Information

**S1:** Compiled dataset.

**S2:** Search results for Web of Science search.

**S3:** Search results for Google Scholar search.

## References

Belote, R. T., & Wilson, M. B. (2020). Delineating greater ecosystems around protected areas to guide conservation. Conservation Science and Practice, 2(6). 10.1111/csp2.196

Brockington, D. (2004). Community conservation, inequality and injustice: myths of power in protected area management. Conservation and Society, 411–432.

Butt, N., Whiting, S., & Dethmers, K. (2016). Identifying future sea turtle conservation areas under climate change. Biological Conservation, 204, 189–196. 10.1016/j.biocon.2016.10.012

Chen, C., Brodie, J. F., Kays, R., Davies, T. J., Liu, R., Fisher, J. T., Ahumada, J., McShea, W., Sheil, D., Agwanda, B., Andrianarisoa, M. H., Appleton, R. D., Bitariho, R., Espinosa, S., Grigione, M. M., Helgen, K. M., Hubbard, A., Hurtado, C. M., Jansen, P. A., … Burton, A. C. (2022). Global camera trap synthesis highlights the importance of protected areas in maintaining mammal diversity. Conservation Letters, 15(2). 10.1111/conl.12865

Climate Change 2021: The physical science basis. Contribution of Working Group I to the Sixth Assessment Report of the Intergovernmental Panel on Climate Change. (2021).

Easterling, D. R., Meehl, G. A., Parmesan, C., Changnon, S. A., Karl, T. R., & Mearns, L. O. (2000). Climate Extremes: Observations, Modeling, and Impacts. Science, 289(5487), 2068–2074. 10.1126/science.289.5487.2068

Field, C. B., Barros, V., Stocker, T. F., Dahe, Q., Dokken, D. J., Plattner, G.-K., Ebi, K. L., Allen, S. K., Mastandrea, M. D., Tignor, M., Mach, K. J., & Midgeley, P. M. (2012). Managing the Risks of Extreme Events and Disasters to Advance Climate Change Adaptation.

Fogarty, D. T., Roberts, C. P., Uden, D. R., Donovan, V. M., Allen, C. R., Naugle, D. E., Jones, M. O., Allred, B. W., & Twidwell, D. (2020). Woody Plant Encroachment and the Sustainability of Priority Conservation Areas. Sustainability, 12(20), 8321. 10.3390/su12208321

Gillingham, P. K., Bradbury, R. B., Roy, D. B., Anderson, B. J., Baxter, J. M., Bourn, N. A. D., Crick, H. Q. P., Findon, R. A., Fox, R., Franco, A., Hill, J. K., Hodgson, J. A., Holt, A. R., Morecroft, M. D., O’Hanlon, N. J., Oliver, T. H., Pearce-Higgins, J. W., Procter, D. A., Thomas, J. A., … Thomas, C. D. (2015). The effectiveness of protected areas in the conservation of species with changing geographical ranges. Biological Journal of the Linnean Society, 115(3), 707–717. 10.1111/bij.12506

Guidelines for Applying Protected Area Management Categories. (2008).

Hannah, L., Midgley, G., Andelman, S., Araújo, M., Hughes, G., Martinez-Meyer, E., Pearson, R., & Williams, P. (2007). Protected area needs in a changing climate. Frontiers in Ecology and the Environment, 5(3), 131–138. 10.1890/1540-9295(2007)5[131:PANIAC]2.0.CO;2

Kunming-Montreal Global Biodiversity Framework (15; L.25). (2022).

Le Saout, S., Hoffmann, M., Shi, Y., Hughes, A., Bernard, C., Brooks, T. M., Bertzky, B., Butchart, S. H. M., Stuart, S. N., Badman, T., & Rodrigues, A. S. L. (2013). Protected Areas and Effective Biodiversity Conservation. Science, 342(6160), 803–805. 10.1126/science.1239268

Leroux, S. J., Schmiegelow, F. K. A., Lessard, R. B., & Cumming, S. G. (2007). Minimum dynamic reserves: A framework for determining reserve size in ecosystems structured by large disturbances. Biological Conservation, 138(3–4), 464–473. 10.1016/j.biocon.2007.05.012

Lewis, S. C., & King, A. D. (2017). Evolution of mean, variance and extremes in 21st century temperatures. Weather and Climate Extremes, 15, 1–10. 10.1016/j.wace.2016.11.002

Loarie, S. R., Duffy, P. B., Hamilton, H., Asner, G. P., Field, C. B., & Ackerly, D. D. (2009). The velocity of climate change. Nature, 462(7276), 1052–1055. 10.1038/nature08649

Maxwell, S. L., Butt, N., Maron, M., McAlpine, C. A., Chapman, S., Ullmann, A., Segan, D. B., & Watson, J. E. M. (2019). Conservation implications of ecological responses to extreme weather and climate events. Diversity and Distributions, 25(4), 613–625. 10.1111/ddi.12878

Mearns, L. O., Watz, R. K., & Schneider, S. H. (1984). Extreme High-Temperature Events: Changes in their probabilities with Changes in Mean Temperature. Journal of Applied Meteorology and Climatology, 23(12), 1601–1613.

Menzel, A., Sparks, T. H., Estrella, N., & Roy, D. B. (2006). Altered geographic and temporal variability in phenology in response to climate change. Global Ecology and Biogeography, 15(5), 498–504. 10.1111/j.1466-822X.2006.00247.x

Nagendra, H. (2008). Do Parks Work? Impact of Protected Areas on Land Cover Clearing. AMBIO: A Journal of the Human Environment, 37(5), 330–337. 10.1579/06-R-184.1

Naughton-Treves, L., Holland, M. B., & Brandon, K. (2005). THE ROLE OF PROTECTED AREAS IN CONSERVING BIODIVERSITY AND SUSTAINING LOCAL LIVELIHOODS. Annual Review of Environment and Resources, 30(1), 219–252. 10.1146/annurev.energy.30.050504.164507

Oldekop, J. A., Holmes, G., Harris, W. E., & Evans, K. L. (2016). A global assessment of the social and conservation outcomes of protected areas. Conservation Biology, 30(1), 133–141. 10.1111/cobi.12568

Parey, S., Dacunha-Castelle, D., & Hoang, T. T. H. (2010). Mean and variance evolutions of the hot and cold temperatures in Europe. Climate Dynamics, 34(2–3), 345–359. 10.1007/s00382-009-0557-0

Pressey, R. L., Cabeza, M., Watts, M. E., Cowling, R. M., & Wilson, K. A. (2007). Conservation planning in a changing world. Trends in Ecology & Evolution, 22(11), 583–592. 10.1016/j.tree.2007.10.001

Roche, D. G., O’Dea, R. E., Kerr, K. A., Rytwinski, T., Schuster, R., Nguyen, V. M., Young, N., Bennett, J. R., & Cooke, S. J. (2022). Closing the knowledge□action gap in conservation with open science. Conservation Biology, 36(3). 10.1111/cobi.13835

Rodrigues, A. S. L., Andelman, S. J., Bakarr, M. I., Boitani, L., Brooks, T. M., Cowling, R. M., Fishpool, L. D. C., da Fonseca, G. A. B., Gaston, K. J., Hoffmann, M., Long, J. S., Marquet, P. A., Pilgrim, J. D., Pressey, R. L., Schipper, J., Sechrest, W., Stuart, S. N., Underhill, L. G., Waller, R. W., … Yan, X. (2004). Effectiveness of the global protected area network in representing species diversity. Nature, 428(6983), 640–643. 10.1038/nature02422

Silver, A., & Andrey, J. (2019). Public attention to extreme weather as reflected by social media activity. Journal of Contingencies and Crisis Management, 27(4), 346–358. 10.1111/1468-5973.12265

Stanton, J. C., Pearson, R. G., Horning, N., Ersts, P., & Reşit Akçakaya, H. (2012). Combining static and dynamic variables in species distribution models under climate change. Methods in Ecology and Evolution, 3(2), 349–357. 10.1111/j.2041-210X.2011.00157.x

Stott, P. (2016). How climate change affects extreme weather events. Science, 352(6293), 1517–1518. 10.1126/science.aaf7271

Stouffer, R. J., & Wetherald, R. T. (2007). Changes of Variability in Response to Increasing Greenhouse Gases. Part I: Temperature. Journal of Climate, 20(21), 5455–5467. 10.1175/2007JCLI1384.1

Sutton, L. J., Ibañez, J. C., Salvador, D. I., Taraya, R. L., Opiso, G. S., Senarillos, T. L. P., & McClure, C. J. W. (2023). Priority conservation areas and a global population estimate for the critically endangered Philippine Eagle. Animal Conservation, 26(5), 684–700. 10.1111/acv.12854

Thompson, R. M., Beardall, J., Beringer, J., Grace, M., & Sardina, P. (2013). Means and extremes: building variability into community□level climate change experiments. Ecology Letters, 16(6), 799–806. 10.1111/ele.12095

Thornton, P. K., Ericksen, P. J., Herrero, M., & Challinor, A. J. (2014). Climate variability and vulnerability to climate change: a review. Global Change Biology, 20(11), 3313–3328. 10.1111/gcb.12581

Trenberth, K. E., Fasullo, J. T., & Shepherd, T. G. (2015). Attribution of climate extreme events. Nature Climate Change, 5(8), 725–730. 10.1038/nclimate2657

Vincenzi, S. (2014). Extinction risk and eco-evolutionary dynamics in a variable environment with increasing frequency of extreme events. Journal of The Royal Society Interface, 11(97), 20140441. 10.1098/rsif.2014.0441

White, H. J., Caplat, P., Emmerson, M. C., & Yearsley, J. M. (2021). Predicting future stability of ecosystem functioning under climate change. Agriculture, Ecosystems & Environment, 320, 107600. 10.1016/j.agee.2021.107600

Zeng, Y., Koh, L. P., & Wilcove, D. S. (2022). Gains in biodiversity conservation and ecosystem services from the expansion of the planet’s protected areas. Science Advances, 8(22). 10.1126/sciadv.abl9885

